# A neural modeling approach to study mechanisms underlying the heterogeneity of visual spatial frequency sensitivity in schizophrenia

**DOI:** 10.1101/2023.10.18.563001

**Authors:** Caroline Dugan, Basilis Zikopoulos, Arash Yazdanbakhsh

**Author notes:** **Correspondence:** Corresponding Authors, Arash Yazdanbakhsh (64 Cummington Mall, Department of Psychological & Brain Sciences, Boston University, MA, 02215); Basilis Zikopoulos (635 Commonwealth Avenue, Department of Health Sciences Boston University, and Department of Anatomy & Neurobiology, Boston University School of Medicine, Boston MA 02215). **Funding:** NIMH R01 MH118500 to BZ and AY.

## Abstract

Patients with schizophrenia exhibit abnormalities in spatial frequency sensitivity, and it is believed that these abnormalities indicate more widespread dysfunction and dysregulation of bottom-up processing. The early visual system, including the first-order Lateral Geniculate Nucleus of the thalamus (LGN) and the primary visual cortex (V1), are key contributors to spatial frequency sensitivity. Medicated and unmedicated patients with schizophrenia exhibit contrasting changes in spatial frequency sensitivity, thus making it a useful probe for examining potential effects of the disorder and antipsychotic medications in neural processing. We constructed a parameterized, rate-based neural model of on-center/off-surround neurons in the early visual system to investigate the impacts of changes to the excitatory and inhibitory receptive field subfields. By incorporating changes in both the excitatory and inhibitory subfields that are associated with pathophysiological findings in schizophrenia, the model successfully replicated perceptual data from behavioral/functional studies involving medicated and unmedicated patients. Among several plausible mechanisms, our results highlight the dampening of excitation and/or increase in the spread and strength of the inhibitory subfield in medicated patients and the contrasting decreased spread and strength of inhibition in unmedicated patients. Given that the model was successful at replicating results from perceptual data under a variety of conditions, these elements of the receptive field may be useful markers for the imbalances seen in patients with schizophrenia.

## Introduction

Schizophrenia is characterized by dysfunction in sensory processing that alters sensory perception, but is distinct from hallucinations [1]. Spatial frequency sensitivity, which in the visual domain describes sensitivity to patterns composed of alternating light and dark bars in a given unit of space, typically in one degree of visual angle, is a central process that is affected in schizophrenia (reviewed in [2]). Spatial frequency is expressed as cycles per degree of sine-wave gratings [3]. While spatial frequency varies to a certain extent between individuals, people tend to be best at detecting light-dark contrast at intermediate spatial frequencies. A variety of factors can affect one’s spatial frequency sensitivity, and abnormal spatial frequency sensitivity patterns have been identified in both medicated and unmedicated patients with schizophrenia.

Medicated patients with schizophrenia who are taking either typical or atypical antipsychotics tend to exhibit decreased contrast sensitivity, although there is conflicting data regarding the range of spatial frequencies at which deficits are observed. Previous studies have demonstrated that medicated patients can exhibit decreased sensitivity across all spatial frequencies [4–6] but in some cases only at low [3, 7–9] or only at medium to high spatial frequencies [9, 10] (Table 1). Factors such as illness duration, medication type, and symptom type have been demonstrated to affect spatial frequency sensitivity, but a general trend of decreased sensitivity in medicated patients is consistent across studies, and it appears that this trend may be more pronounced with increased illness duration. Studies indicate that patients with an illness duration greater than ten years have decreased contrast sensitivity at all spatial frequencies [6], whereas medicated patients with an illness duration less than ten years exhibit decreased sensitivity at only low spatial frequencies [9]. With regards to medication type, patients taking atypical antipsychotics may experience less pronounced deficits in contrast sensitivity compared with patients taking typical antipsychotics [6, 9].

**Table 1.**
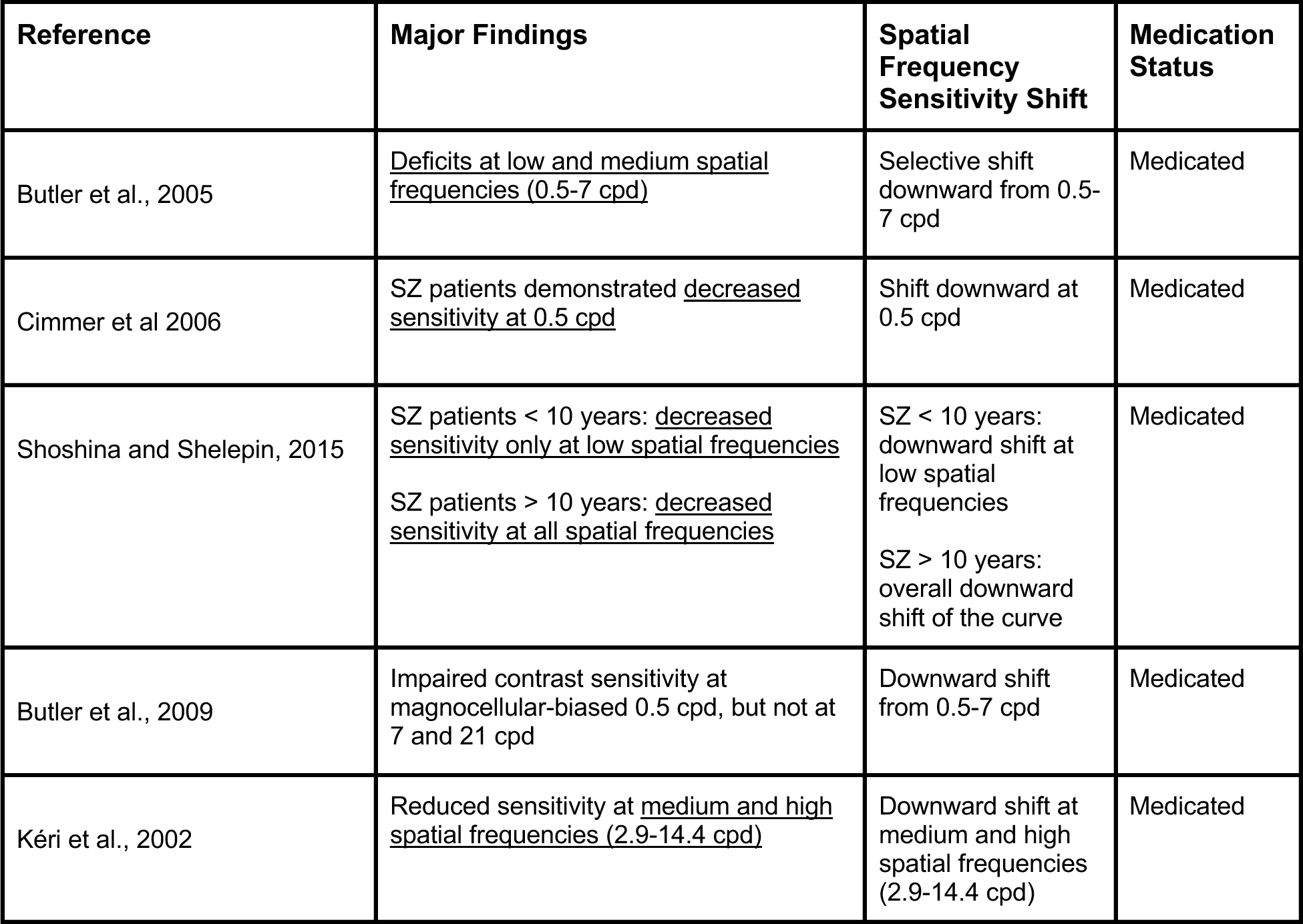
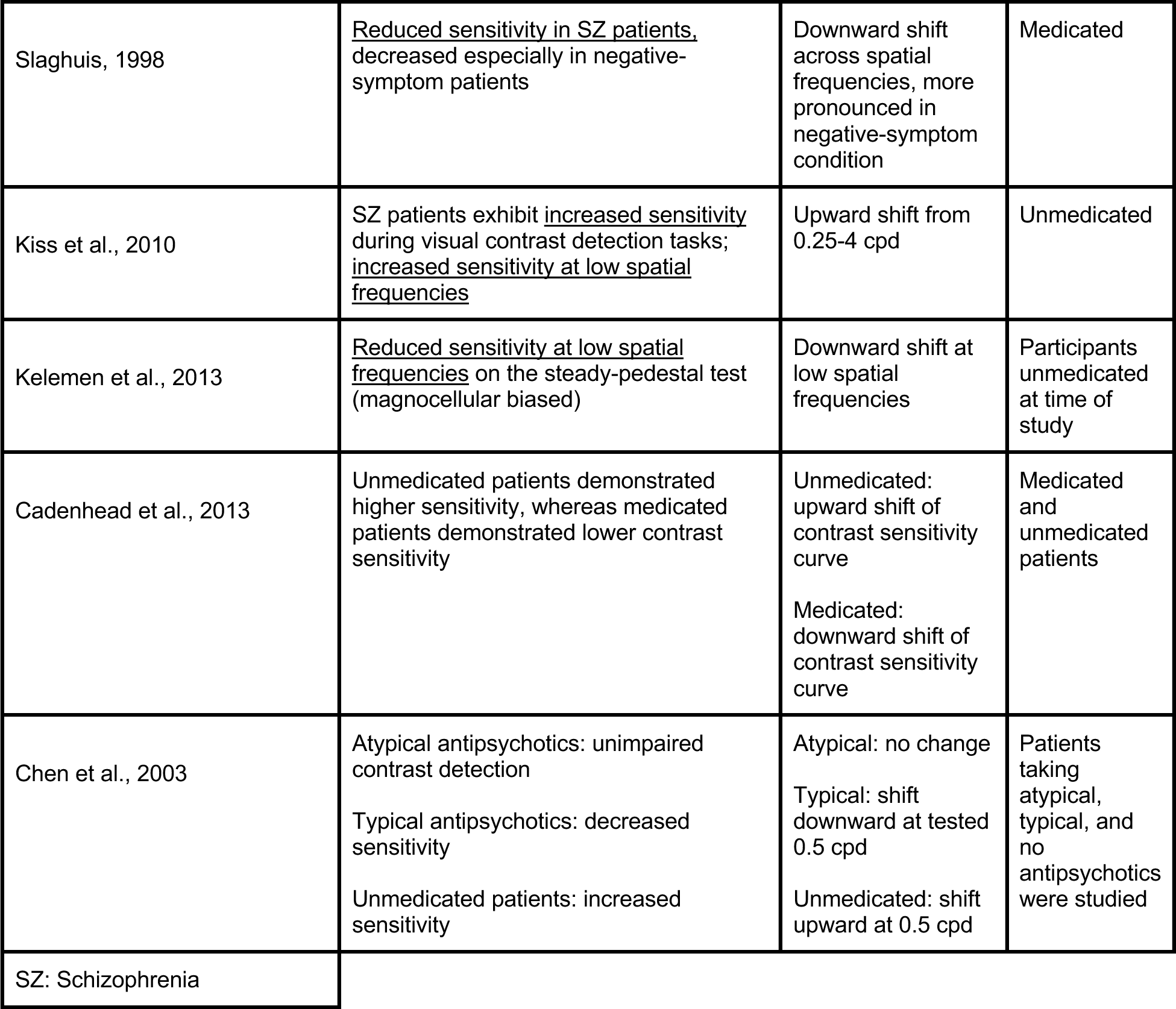
Relevant findings from previous studies of spatial frequency sensitivity in patients with schizophrenia that we replicated with the single-layer feedforward visual spatial frequency sensitivity model.

| Reference | Major Findings | Spatial Frequency Sensitivity Shift | Medication Status |
| --- | --- | --- | --- |
| Butler et al., 2005 | <u>Deficits at low and medium spatial frequencies (0.5-7 cpd)</u> | Selective shift downward from 0.5-7 cpd | Medicated |
| Cimmer et al 2006 | SZ patients demonstrated <u>decreased sensitivity at 0.5 cpd</u> | Shift downward at 0.5 cpd | Medicated |
| Shoshina and Shelepin, 2015 | SZ patients < 10 years: <u>decreased sensitivity only at low spatial frequencies</u><br>SZ patients > 10 years: <u>decreased sensitivity at all spatial frequencies</u> | SZ < 10 years: downward shift at low spatial frequencies<br>SZ > 10 years: overall downward shift of the curve | Medicated |
| Butler et al., 2009 | Impaired contrast sensitivity at magnocellular-biased 0.5 cpd, but not at 7 and 21 cpd | Downward shift from 0.5-7 cpd | Medicated |
| Kéri et al., 2002 | Reduced sensitivity at <u>medium and high spatial frequencies (2.9-14.4 cpd)</u> | Downward shift at medium and high spatial frequencies (2.9-14.4 cpd) | Medicated |
| Slaghuis, 1998 | <u>Reduced sensitivity in SZ patients</u>, decreased especially in negative-symptom patients | Downward shift across spatial frequencies, more pronounced in negative-symptom condition | Medicated |
| Kiss et al., 2010 | SZ patients exhibit <u>increased sensitivity</u> during visual contrast detection tasks; <u>increased sensitivity at low spatial frequencies</u> | Upward shift from 0.25-4 cpd | Unmedicated |
| Kelemen et al., 2013 | <u>Reduced sensitivity at low spatial frequencies</u> on the steady-pedestal test (magnocellular biased) | Downward shift at low spatial frequencies | Participants unmedicated at time of study |
| Cadenhead et al., 2013 | Unmedicated patients demonstrated higher sensitivity, whereas medicated patients demonstrated lower contrast sensitivity | Unmedicated: upward shift of contrast sensitivity curve<br>Medicated: downward shift of contrast sensitivity curve | Medicated and unmedicated patients |
| Chen et al., 2003 | Atypical antipsychotics: unimpaired contrast detection<br>Typical antipsychotics: decreased sensitivity<br>Unmedicated patients: increased sensitivity | Atypical: no change<br>Typical: shift downward at tested 0.5 cpd<br>Unmedicated: shift upward at 0.5 cpd | Patients taking atypical, typical, and no antipsychotics were studied |
| SZ: Schizophrenia |  |  |  |

Despite challenges in research involving unmedicated patients with schizophrenia, studies have identified that, in general, unmedicated patients tend to exhibit increased contrast sensitivity. Some observed this increase at low spatial frequencies [11, 12], while others found increased sensitivity across all spatial frequencies (Table 1) [5]. The spatial frequency range at which abnormalities are observed may be dependent on illness stage, as first-episode, medication-naïve patients have been shown to exhibit increased sensitivity at low spatial frequencies [12].

In addition to visual perception changes and hallucinations, visual processing abnormalities have been linked to several aspects of schizophrenia. Certain visual changes in at-risk individuals have been associated with later development of schizophrenia, and visual impairments are correlated with reduced real-world functioning in patients [13–16]. Additionally, low-level visual processing abnormalities in schizophrenia, such as altered spatial frequency sensitivity, may indicate more widespread dysfunction, suggesting that there is a bottom-up component of the disorder [17]. Growing evidence suggests that deficits in sensory processing in individuals with schizophrenia can provide clues into the disorder’s overall mechanism. However, even though the visual system provides distinct advantages for the study of sensory processes [18], the involvement of visual circuit components and mechanisms in the pathophysiology of schizophrenia is not well understood. To address this gap, we developed a rate-based feedforward model to test the overall impact of hypothesized changes in bottom-up visual excitatory networks that take into account local inhibitory circuit modulation. We investigated the model’s performance to illustrate how sensory processing abnormalities in schizophrenia connect perceptual deficits with potential underlying excitatory and inhibitory changes. By modulating the excitatory and inhibitory receptive field subfields, we translated all receptive field changes to deviations in the balance of excitation and inhibition; from there, we were able to investigate the perceptual outcome and connect the neural modeling results to previous data collected in perceptual studies.

## Materials and Methods

We developed a rate-based neural model of a primary visual circuit that can simulate bottom-up processing of pairs of interconnected areas in the central visual pathway, including a) retinal input to the lateral geniculate nucleus (LGN) of the thalamus, the first-order visual thalamic nucleus that relays visual input to the cortex, b) from LGN to the primary visual cortex (area 17), and c) feedforward corticocortical connections between visual association areas [19–27]. The parameterized, rate-based, feedforward neural model (Equation 1), which was generated using MATLAB (2019b), receives an input array, and through its connections excites and inhibits the 4000 model neurons. The connectivity follows an on-center/off-surround organization [28–37]. The current model size allows for sampling of visual inputs with the appropriate resolution. Receptive field organization was modeled using a Difference of Gaussians (DoG) method. Model neurons were then presented with sinusoidal grating input with varying spatial frequencies, and the model’s response was measured to generate contrast sensitivity curves (Figure 1A). Following a broad range of single or combined parameter changes, described in detail below, we compared model responses with key behavioral/perceptual findings from previous studies of spatial frequency sensitivity in patients with schizophrenia (Table 1), with the aim to identify possible circuit mechanisms and changes at the receptor or neurotransmitter levels at multiple states of the disorder.

**Figure 1.**
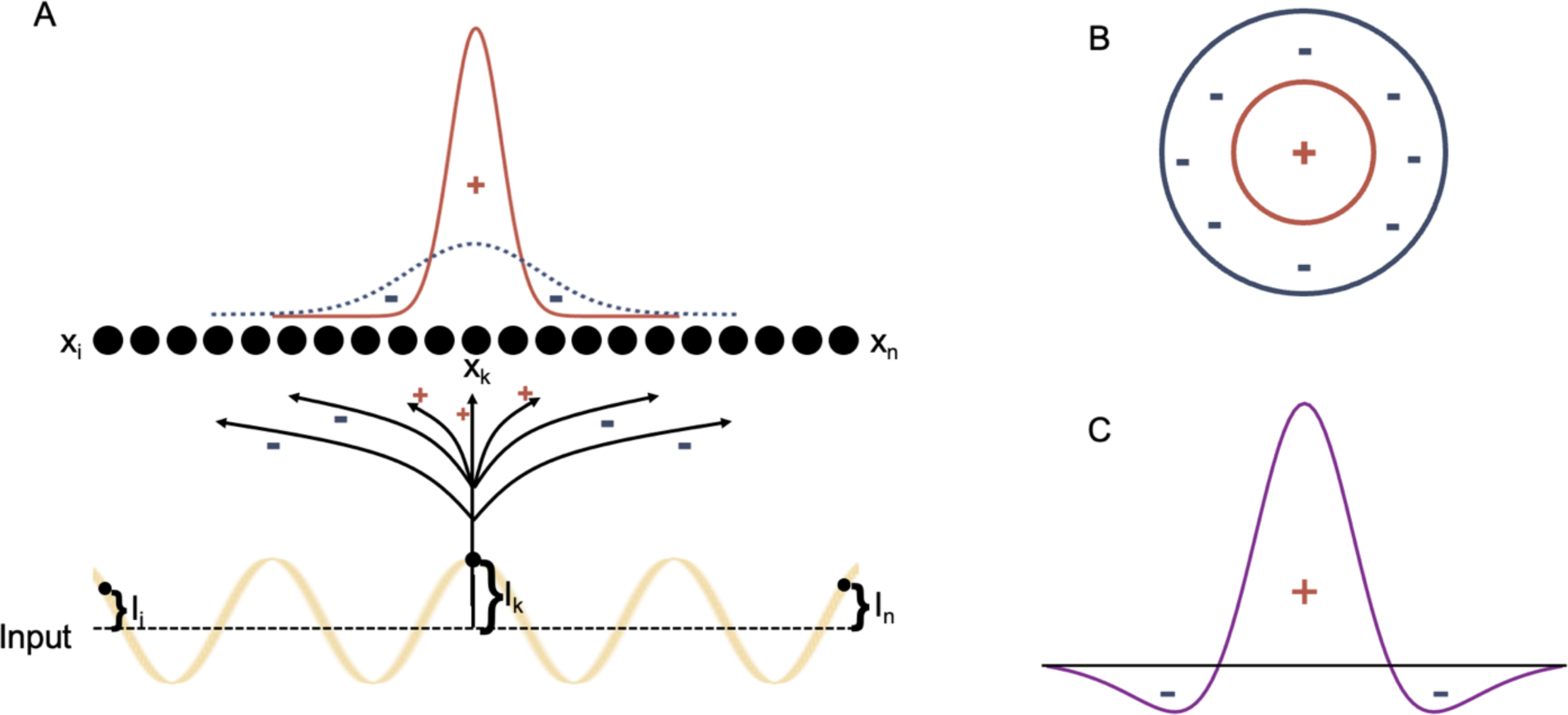
Model Network and Receptive Field Model. Array of model neurons receiving input through excitatory and inhibitory connections. The studied inputs are luminance sin modulation (A). Considering an on-center/off-surround organization of receptive fields (B), in 1D, we implemented excitatory and inhibitory connectivities following weighted subtraction of Gaussians (Difference of Gaussians, DoG) to approximate excitatory and inhibitory subfields of receptive field.

### Difference of Gaussians

The Difference of Gaussians (DoG) provides an on-center/off-surround parametrizable method for the model receptive fields. The Gaussians for the excitatory on-center and inhibitory off-surround are concentric and the difference between the two approximates an on-center/off-surround receptive field organization. With the model feedforward architecture from the input array to the model neuron array following on-center/off-surround interaction, we measured the model’s response to visual stimuli with varying spatial frequency.

### Neural Activity

The activity of each neuron in the model, represented by vector **x** (Figure 1), is determined by the following shunting equation [32, 33, 38, 39]:

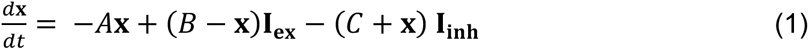

Constants A, B, and C represent the neural activity decay term, neural activity upper limit, and neural activity lower limit, respectively. Bold variables represent arrays. Model neurons array activity (**x**) was dependent on the excitatory (**I_ex_**) and the inhibitory (**I_inh_**) components of the input array: **I_ex_** refers to the input array (**I**) convolved with the excitatory Gaussian (**G_ex_**), and **I_inh_** refers to the input array (**I**) convolved (*) with the inhibitory Gaussian (**G_inh_**):

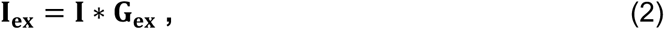

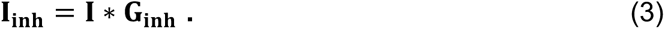

G_ex_ and G_inh_ follow the equations:

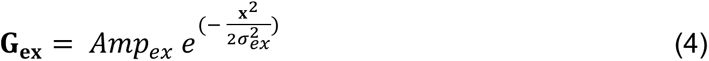

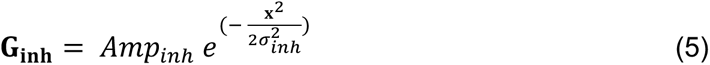

In which **x** ranges from −5 *σ_ex/inh_* to 5 *σ_ex/inh_*. All parameters and symbols are summarized in Table 1 with their values in Supplementary Table 1. Amp_ex_ and Amp_inh_ refer to the amplitude or height of the excitatory and inhibitory subfields (Gaussians), respectively. σ_ex_ and σ_inh_ refer to the width of the excitatory and inhibitory subfields (Gaussians), respectively.

Our current neural modeling results are based on the equilibrium state of neural dynamics of Equation 1:

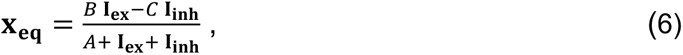

where **x_eq_** is model neurons array activity at equilibrium.

Substituting **I_ex_** and **I_inh_** values from Eqs. 2-3 yields:

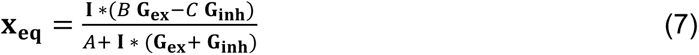

The equilibrium of Eq. 7 shows the normalized by input weighted subtraction of excitatory and inhibitory Gaussian or Difference of Gaussians (DoG) as the effective combined excitatory and inhibitory subfields processing the model inputs.

Considering DoG, we set the inhibitory Gaussian, with a larger σ (σ_inh_) and smaller amplitude (Amp_inh_), and the excitatory Gaussian, with a smaller sigma (σ_ex_) and a larger amplitude (Amp_ex_) for all of the reported results. The result of such Gaussians’ subtraction mimics an on-center/off-surround receptive field [40], given that it has a positive center flanked by two negative, inhibitory regions (Figure 1). Marr and Hildreth [40] showed that the ratio of σ_inh_ to σ_ex_ for optimal contrast registration is 1:1.6, which we implemented for the base state of simulations and by varying σs, amplitudes, and combinations, we examined their impacts on the neural model spatial frequency sensitivity.

### Input Generation

To determine the model’s response to varying spatial frequencies, sinusoidal inputs with varying spatial frequencies were generated and used (Equation 8). These inputs represent luminance gratings with varying spatial frequencies (Figure 2). The peaks of the sinusoidal gratings correspond to areas of maximum luminance, while the troughs of the sinusoidal gratings correspond to areas of minimum luminance.

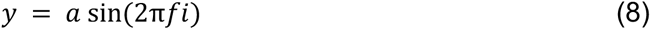

**Figure 2.**
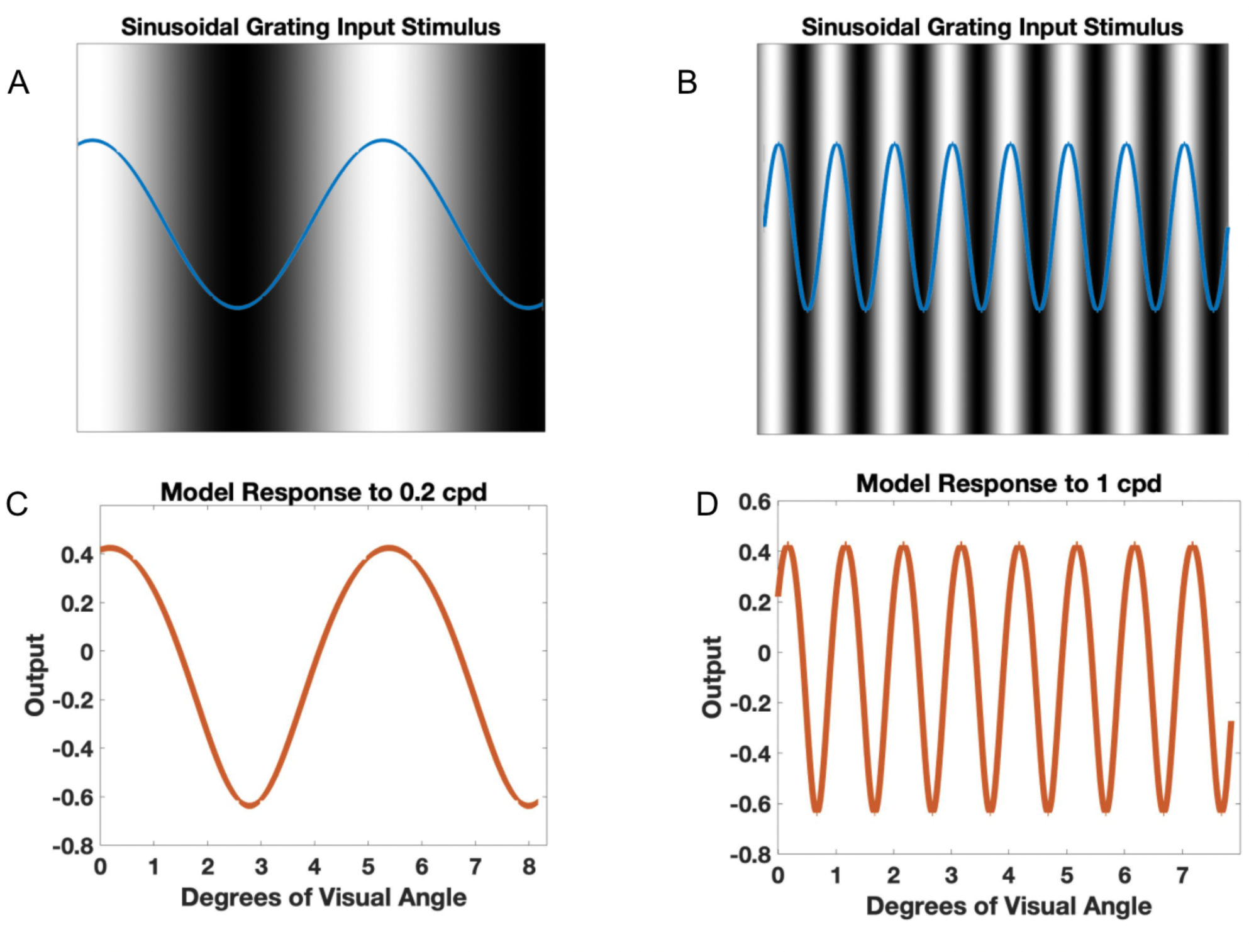
Sinusoidal grating inputs and corresponding model response. Sinusoidal grating inputs of 0.2 (A) and 1 (B) cpd were generated. The blue sin waves represent the input stimulus while the black and white images represent the corresponding luminance gratings they represent, with black areas corresponding to minimum luminance values (trough of sin graph) and white areas corresponding to maximum luminance values (peak of sin graph). The model response to inputs of 0.2 (C) and 1 (D) cpd were then measured.

*a* refers to the amplitude of the sinusoidal grating, which was held constant at 0.1 across model trials. *f* refers to spatial frequency used, which ranged from 0.1 to 100 cycles per degree (cpd). Position is represented by the variable *i*. The model was shown 200 sinusoidal gratings with spatial frequencies ranging from 0.1 to 100 cpd and its response to each grating was measured and plotted to create the contrast sensitivity curves.

### Parameter Selection

Across model trials, amplitude and σ values were altered while values of A, B, and C were kept constant. A, B, and C were set to values of 1, 10.1, and 5. Parameter selections for each model run are reflected in Supplementary Table 1, and Table 2 reflects a summary of the parameters and symbols used in equations.

**Table 2.** Parameters and symbols used in equations.

|  |  |
| --- | --- |
| **Amp<sub>ex</sub>** | Height of the excitatory Gaussian |
| **Amp<sub>inh</sub>** | Height of the inhibitory Gaussian |
| **$\sigma_{ex}$** | Width of the excitatory Gaussian |
| **$\sigma_{inh}$** | Width of the inhibitory Gaussian |
| **\*** | Convolution |
| **A** | Neural activity decay term |
| **B** | Neural activity upper limit |
| **C** | Neural activity lower limit |
| **a** | Sinusoidal amplitude |
| **f** | Spatial frequency |
| **i** | Sinusoidal position |
| **$\theta$** | Cosine similarity angle between sensitivity curves |

### Measuring Model Response

To determine the model’s contrast sensitivity for varying spatial frequencies, the minimum and maximum values of the model neurons array response were determined for each spatial frequency. Given the model’s connectivity, architecture, and the input stimuli used, the model neuron array has maximum activities at maximum luminance, or sinusoidal peak locations, and the minimum activities are at minimum luminance, or sinusoidal trough locations (Figure 2). We therefore consider the contrast readout of the model based on maximum and minimum activities for each sin stimulus as the model representation of contrast sensitivity:

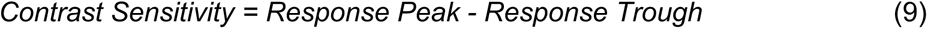

In other words, for each sinusoidal grating input, the model’s representation of contrast sensitivity was determined by subtracting the neuron array’s minimum activity from the maximum activity. Correspondingly, in perceptual studies (e.g., see Table 1), spatial frequency sensitivity is determined by the inverse of observers’ contrast detection threshold at various spatial frequencies as a pointer to the magnitude of neural representation of contrast.

### Spatial Frequency Ranges

By comparing model response patterns with existing data from perceptual studies in patients with schizophrenia, the low, medium, and high spatial frequency ranges were determined [3, 7, 9, 11]. Low spatial frequency ranged between 0.1-4 cpd, medium spatial frequency between 4-10 cpd, and high spatial frequency between 10-25 cpd. We considered consistent ranges for low, mid, and high spatial frequencies, however, the exact ranges in prior studies could vary depending on upper and lower limits for low, medium, and high spatial frequencies as well as the stimulus type (i.e., Gabor vs. uniform spatial frequency). Therefore, certain reported differences from perceptual studies at medium spatial frequencies, for example, may slide into our uniformly defined low spatial frequency range; hence, studies reported in Table 1 should be considered with their uniquely defined spatial frequency ranges.

### Analysis

The model contrast sensitivity is represented with a vector. Cosine similarity (cosSim) based angle (θ) between the model sensitivity vector (MSV) in a given condition and the overall model “base” contrast sensitivity vector (MBCSV) could be calculated using Equations 10 (• stands for dot product) and 11:

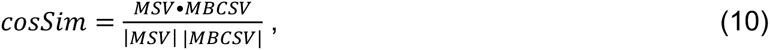

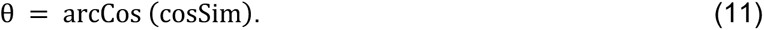

The normalized difference index (NDI) between the MSV and MBCSV at low, medium, and high spatial frequencies was determined according to Equation 12:

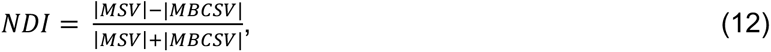

in which |…| represents the vector size, and the NDI significance was determined using t-test.

## Results

By measuring the neural model contrast sensitivity with different excitatory and inhibitory parameters and comparing the model contrast sensitivity with each parameter set with contrast sensitivity perceptual data from medicated and unmedicated patients with schizophrenia as well as healthy controls, we could approximate and compare the excitation / inhibition balance in each group. Supplementary Table 1 summarizes the model’s parameter set modifications from base set (control) that result in matched contrast sensitivity to medicated and unmedicated patients. Table 3 summarizes the model’s parameter set modifications from base set (control) that result in contrast sensitivity changes and the best fit replication that was determined by examining the NDI value and magnitude for each model sensitivity curve at low, medium, and high spatial frequencies as well as the overall curve shape, as reported by cosSim and θ in low, medium, and high spatial frequency ranges (see also Supplementary Table 2).

**Table 3.**
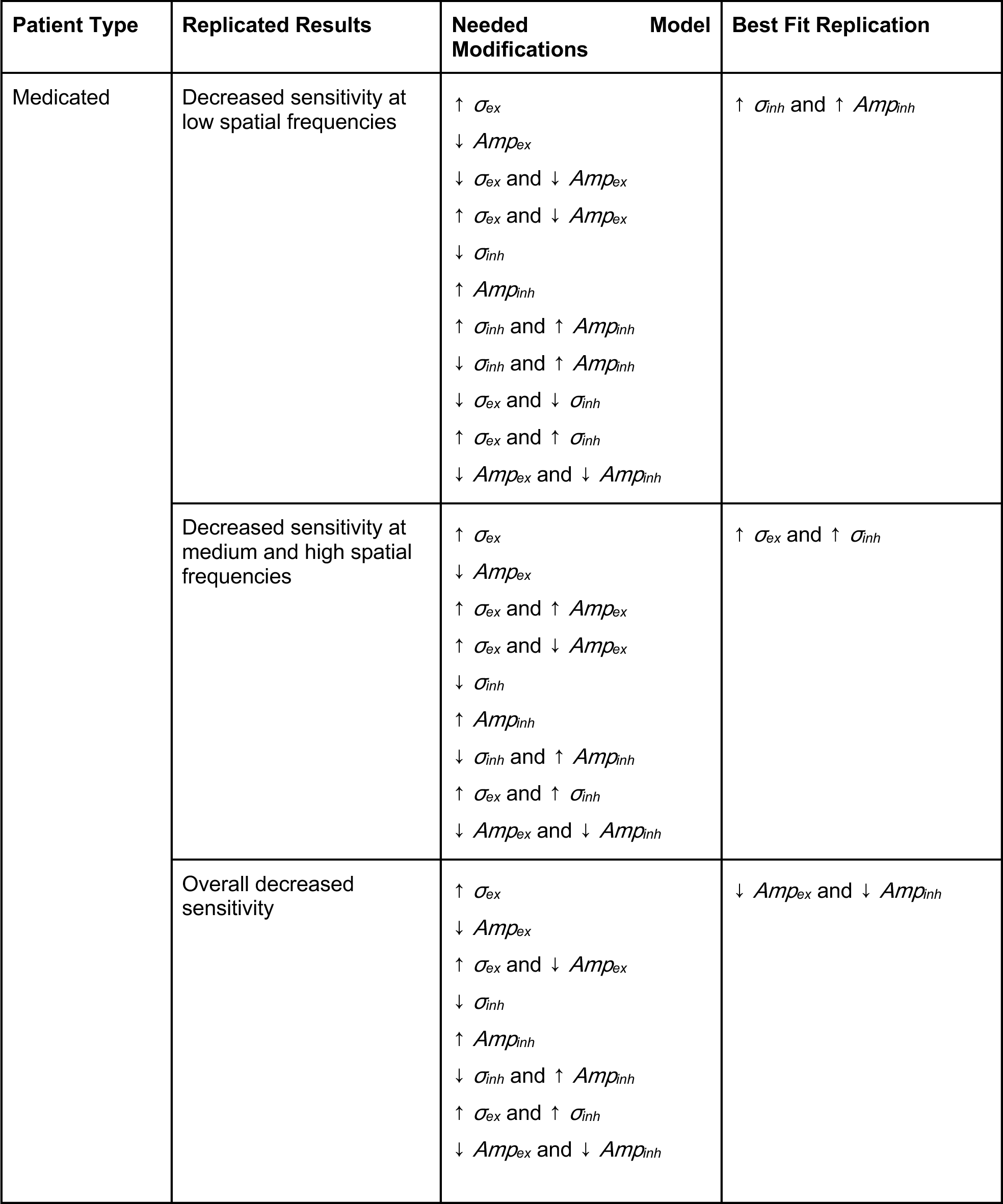
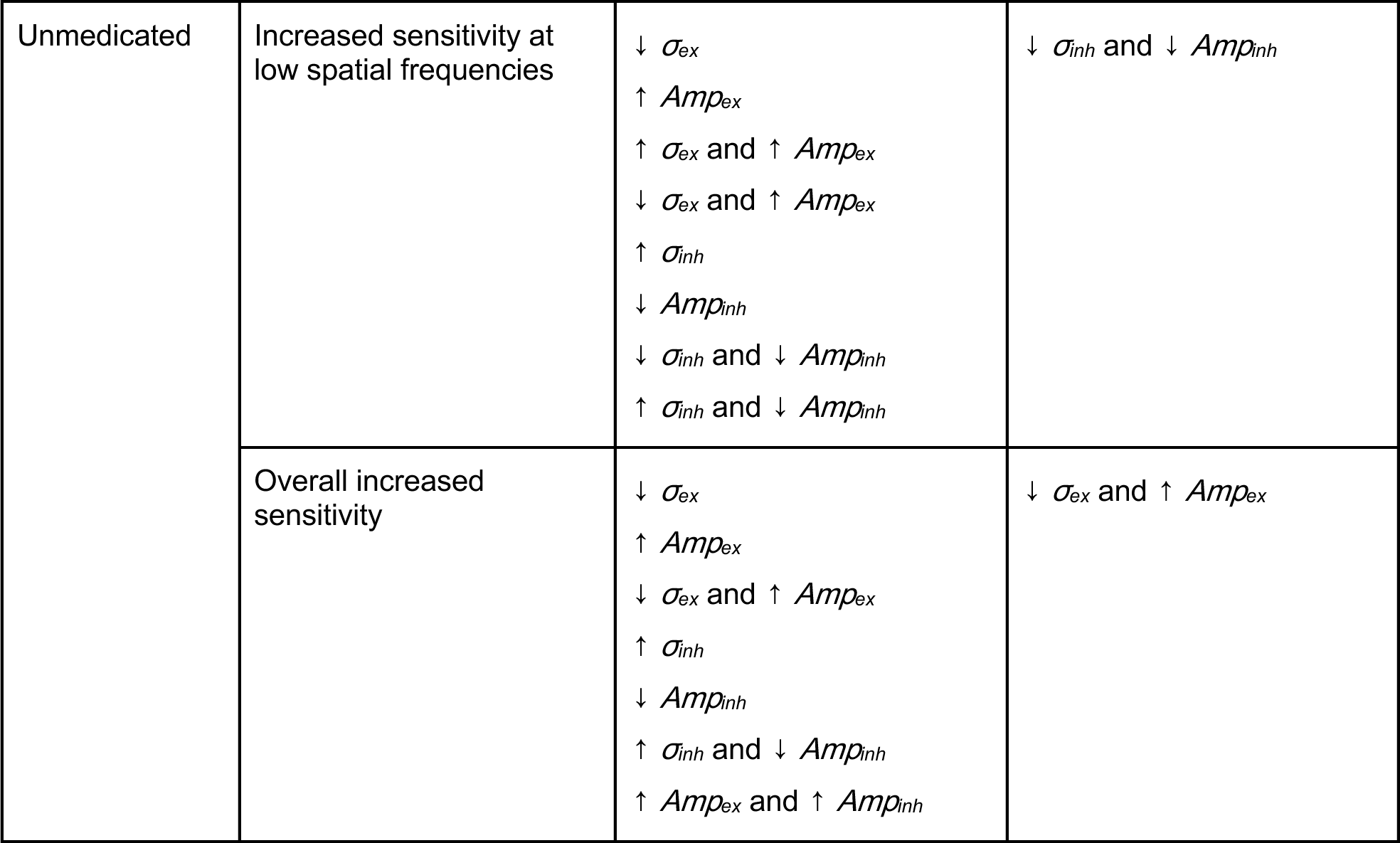
Model excitatory and inhibitory subfield changes matching patient perceptual contrast sensitivity data. Best fit replication was determined by examining the NDI value and magnitude for each model sensitivity curve at low, medium, and high spatial frequencies as well as the overall curve shape, as reported by cosSim and θ (see Supplementary Table 2).

### Effects of Changes in Excitation

#### Impact of Changes in Excitation Extent (σ_ex_)

To determine the isolated impact of altering the excitatory subfield extent (*σ_ex_*), we held *σ_inh_*, *Amp_inh_*, and *Amp_ex_* constant at standard values of 1.6, 1, and 1, respectively. *σ_ex_* was then varied (Supplementary Table 1), and the resulting contrast sensitivity vs. input spatial frequency curves were compared to the control base (Figure 3). Varying *σ_ex_* from the base value of 1 resulted in an overall shift of the contrast sensitivity curve, changing both overall sensitivity and the spatial frequency at which the model was most sensitive. Decreasing *σ_ex_* to a value of 0.8 resulted in increased sensitivity across spatial frequencies, and the spatial frequency at which the model demonstrated highest sensitivity was shifted rightward to higher spatial frequencies when compared with the model base sensitivity curve. Alternatively, increasing *σ_ex_* to a value of 1.2 resulted in decreased sensitivity across spatial frequencies and a shift in maximum sensitivity toward lower spatial frequencies (Figure 3). Therefore, decreasing *σ_ex_* resulted in maximum sensitivity at higher spatial frequencies and overall increased sensitivity, while increasing *σ_ex_* resulted in maximum sensitivity at very low spatial frequencies and overall decreased sensitivity.

**Figure 3.**
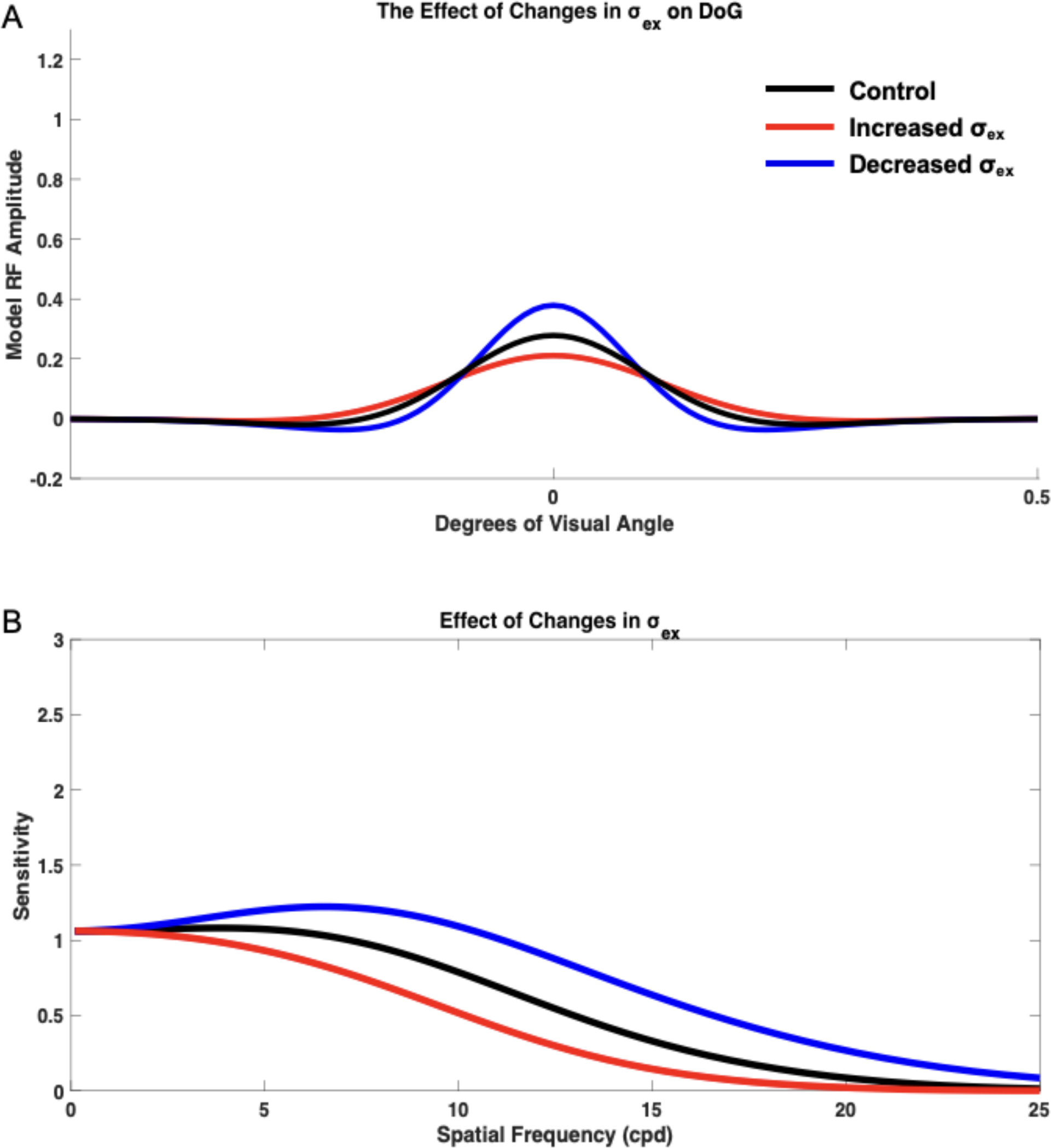
Isolated impact of changing excitatory extent (*σ_ex_*) on the neural model spatial frequency sensitivity and receptive field profile (DoG). DoG profiles with modified excitatory subfield extents (A) and their impacts on the neural model spatial frequency sensitivities (B). Neural model base (control) spatial frequency sensitivity (black), when *σ_ex_* was increased from 1 to 1.2 (red), and decreased to 0.8 (blue).

#### Impact of Changes in Excitation Amplitude (Amp_ex_)

To determine isolated impact of *Amp_ex_*, the three other parameters, *σ_ex_*, *σ_inh_*, and *Amp_inh_* were held constant at standard values of 1, 1.6, and 1, respectively, while *Amp_ex_* was varied (Figure 4). Varying *Amp_ex_* from the base (control) value of 1 resulted in changes in both overall sensitivity and the spatial frequency at which maximum sensitivity occurred. Increasing *Amp_ex_* to a value of 1.2 resulted in an overall increase in sensitivity across spatial frequencies, and the spatial frequency at which the model demonstrated maximum sensitivity was shifted leftward toward lower spatial frequencies when compared with the control. Increasing *Amp_ex_* also resulted in a lack of a distinct peak, with maximum sensitivity occurring within a small range of spatial frequencies rather than at one distinct spatial frequency. Decreasing *Amp_ex_* to 0.8 resulted in an overall decrease in sensitivity across spatial frequencies, and maximum sensitivity occurred at higher spatial frequencies when compared with the base control (*Amp_ex_* = 1). It is also important to note that decreasing *Amp_ex_* resulted in the appearance of a more distinct peak in the spatial frequency sensitivity curve. Thus, varying *Amp_ex_* affected overall sensitivity, the spatial frequency at which maximum sensitivity occurred, and peak emergence in the sensitivity curve.

**Figure 4.**
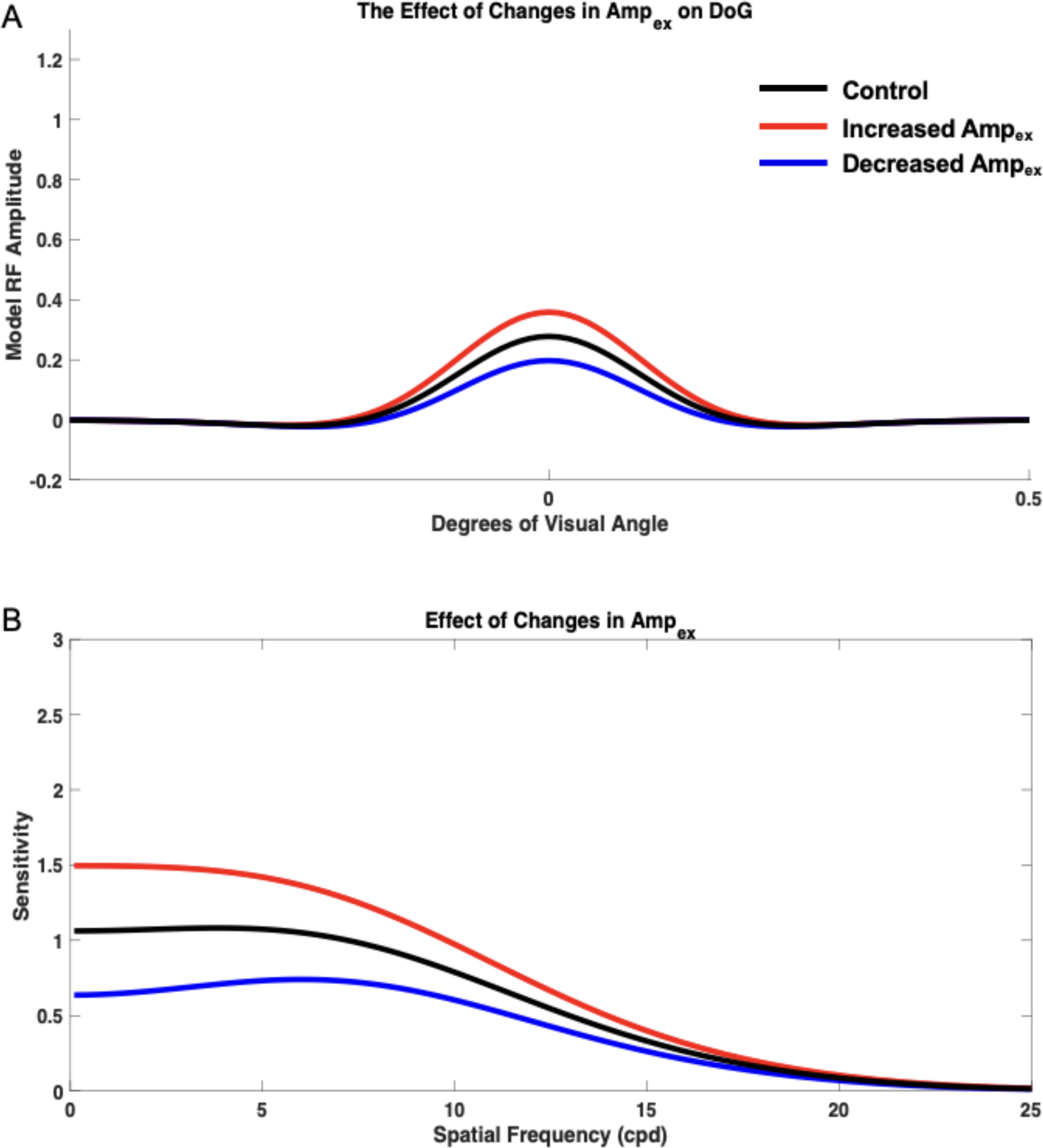
Isolated impact of changing excitatory amplitude (*Amp_ex_*) on the neural model spatial frequency sensitivity and receptive field profile (DoG). DoG profiles with modified excitatory subfield amplitude (A) and their impacts on the neural model spatial frequency sensitivities (B) was compared with the control (black) when *Amp_ex_* was increased to a value of 1.2 (red) and decreased to a value of 0.8 (blue).

#### Impact of Concurrent Changes in Excitatory Spatial Extent (σ_ex_) and Strength (Amp_ex_)

To measure the concurrent impact of *σ_ex_* and *Amp_ex_*, *σ_inh_* and *Amp_inh_* values were held constant at 1.6 and 1, respectively. *Amp_ex_* and *σ_ex_* were then simultaneously varied from their control values of 1 and the resulting contrast sensitivity curves were compared with the control curve (Figure 5). When *σ_ex_* was decreased to 0.6 and *Amp_ex_* was also decreased to 0.6, the model exhibited decreased sensitivity at low spatial frequencies and increased sensitivity at higher spatial frequencies (Figure 5 A-B). Peak sensitivity also shifted to a higher spatial frequency when compared with the control. Increasing *σ_ex_* to 1.4 and *Amp_ex_* to 1.4 resulted in increased sensitivity at low spatial frequencies, into the low end of the medium spatial frequency range, and decreased sensitivity at higher spatial frequencies (Figure 5 C-D). The spatial frequency at which the model was maximally sensitive was lower compared to the control, and the sensitivity curve demonstrated less of a distinct peak. When *σ_ex_* was decreased to 0.6 while *Amp_ex_* was simultaneously increased to 1.4, the model demonstrated increased sensitivity across spatial frequencies, and maximum sensitivity occurred at a higher spatial frequency compared to the control (Figure 5 E-F). Increasing *σ_ex_* to 1.4 while decreasing *Amp_ex_* to 0.6 resulted in overall decreased sensitivity across spatial frequencies, and maximum sensitivity occurred at a lower spatial frequency compared to the control (Figure 5 G-H). Overall, concurrent changing of *σ_ex_* and *Amp_ex_* affected the preferred spatial frequency of the model toward lower and higher spatial frequencies.

**Figure 5.**
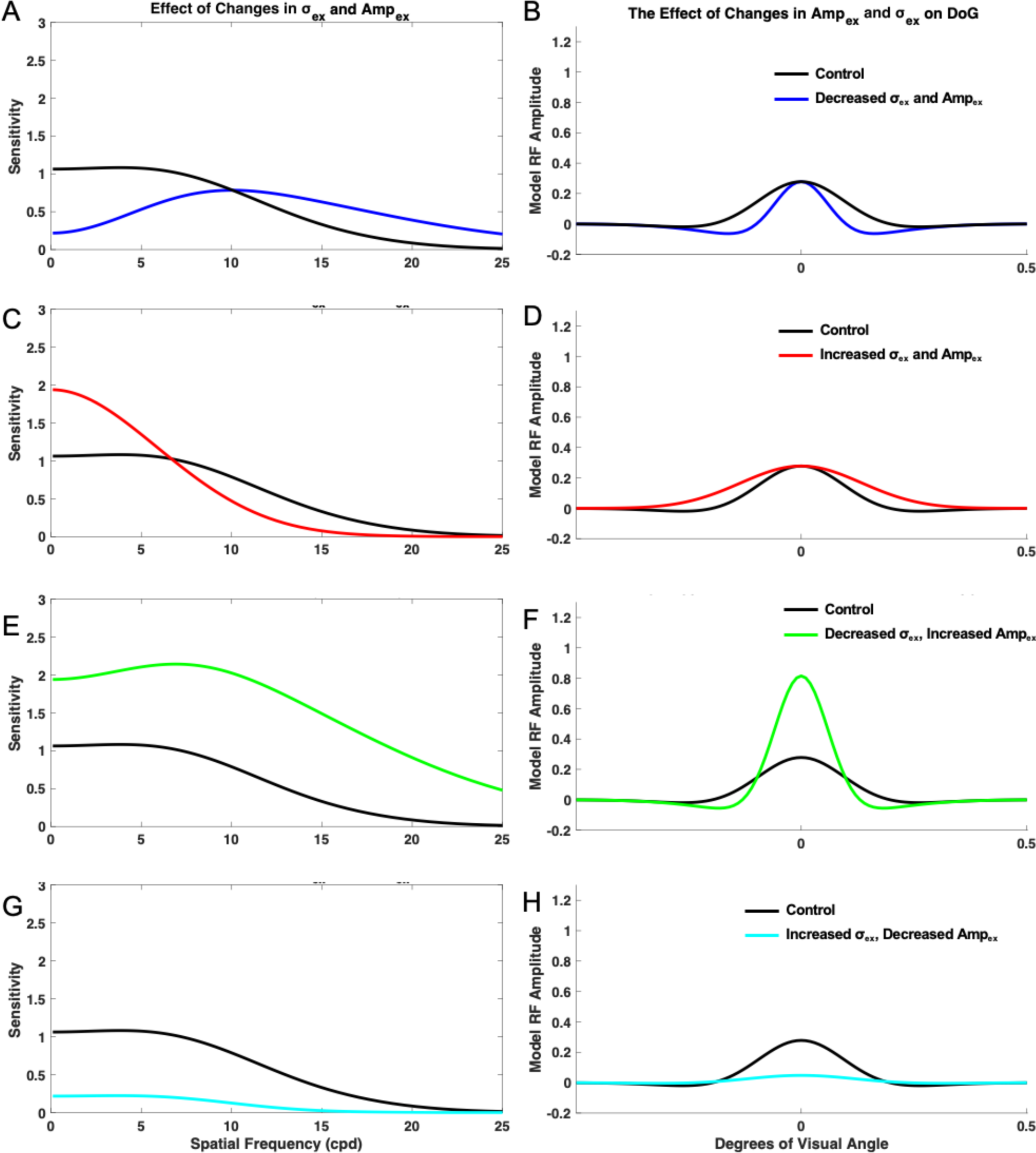
Impact of concurrent changes in excitation extent (*σ_ex_*) and amplitude (*Amp_ex_*) on the neural model receptive field profile (DoG) and spatial frequency sensitivity. Impacts were measured when *σ_ex_* and *Amp_ex_* were decreased to 0.6 (A-B), when *σ_ex_* and *Amp_ex_* were increased to 1.4 (C-D), when *σ_ex_* was decreased to 0.6 and *Amp_ex_* was increased to 1.4 (E-F), and when *σ_ex_* was increased to 1.4 and *Amp_ex_* was decreased to 0.6 (G-H).

### Impacts of Modified Inhibition

#### Impact of Changes in Inhibition Extent (σ_inh_)

To measure the isolated impact of *σ_inh_*, values of *σ_ex_*, *Amp_ex_*, and *Amp_inh_* were held constant at the base value of 1. Then, *σ_inh_* were varied and the resulting contrast sensitivity curves were compared with the control base curve, which was generated using a *σ_inh_* value of 1.6 (Figure 6). Varying *σ_inh_* resulted in changes in overall sensitivity and the peak pattern in the sensitivity curve. Decreasing *σ_inh_* to 1.2 resulted in overall decreased sensitivity, although these effects were more pronounced at low to medium spatial frequencies, with the lack of a distinct peak, although maximum sensitivity occurred at lower spatial frequencies when compared with the base curve. When *σ_inh_* was increased to a value of 2.0, however, the model had a more distinct peak, demonstrating a stronger preference for its peak spatial frequency. The model’s spatial frequency sensitivity curve also demonstrated a slight rightward shift toward higher spatial frequencies. Additionally, the model exhibited increased sensitivity across spatial frequencies, but these effects were more pronounced at low to medium spatial frequencies.

**Figure 6.**
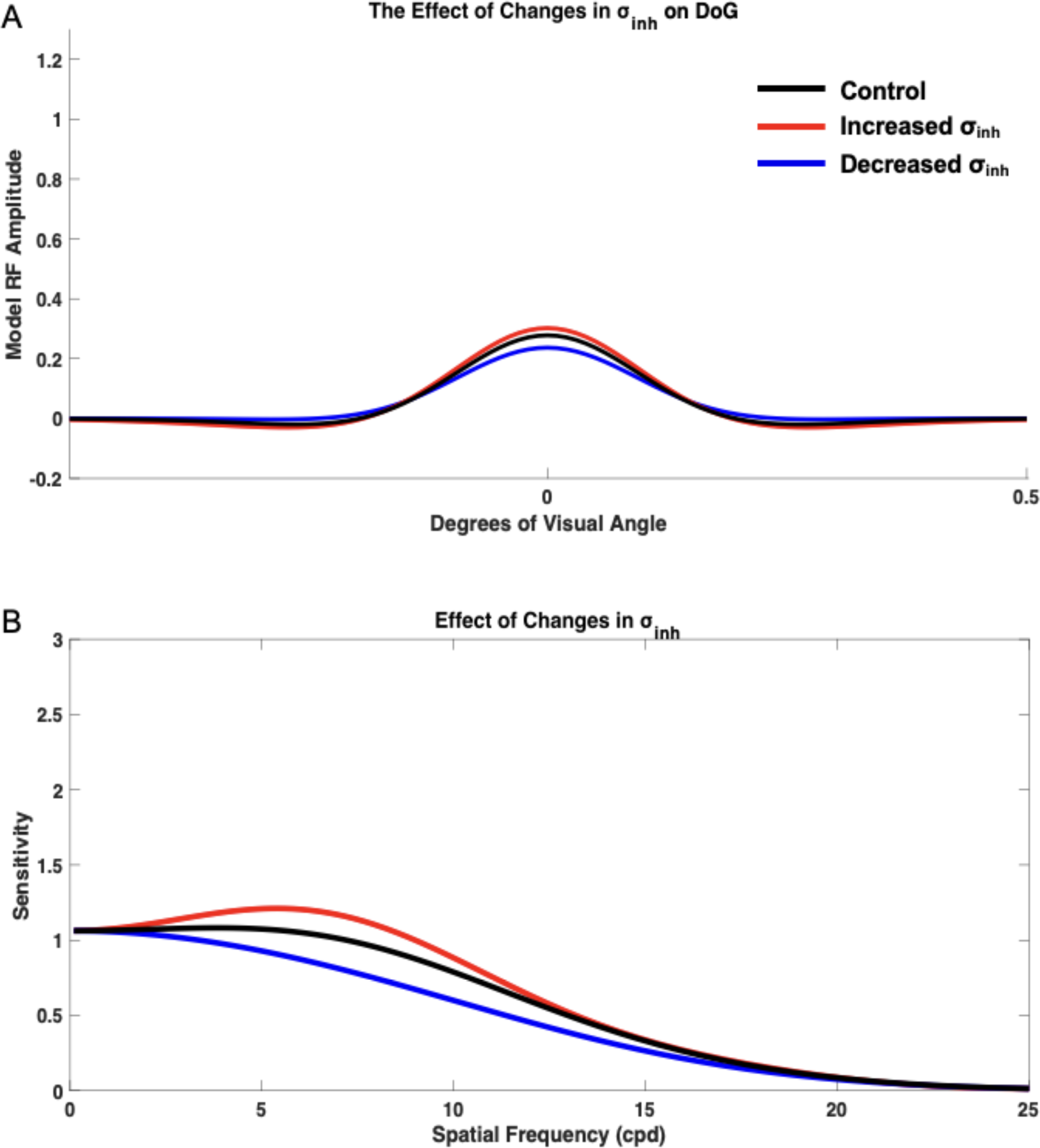
Isolated impact of inhibition extent (*σ_inh_*) on spatial frequency sensitivity and receptive field profile (DoG). DoG profiles (A) and the impact on spatial frequency sensitivity (B) was compared with the control (black) when *σ_inh_* was increased to 2.0 (red) and decreased to 1.2 (blue).

#### Impact of Changes in Inhibition Strength (Amp_inh_)

To measure the isolated impact of *Amp_inh_*, we held σ_inh_, σ_ex_, and Amp_ex_ constant at base values of 1.6, 1, and 1, respectively, and *Amp_inh_* was varied from the base value of 1, and the resulting contrast sensitivity curves were compared with the control curve, which was generated using *Amp_inh_* with value of 1 (Figure 7). Changing *Amp_inh_* most clearly altered the model’s sensitivity at low to medium spatial frequencies as well as the spatial frequency at which the model was maximally sensitive. Decreasing *Amp_inh_* from based 1 to 0.6 (from its base 1) resulted in increased sensitivity across spatial frequencies, although these effects were most pronounced at low to medium spatial frequencies, and a shift in preference toward low spatial frequencies. The model’s response also lacked a distinct peak when *Amp_inh_* was decreased. Increasing *Amp_inh_* from base 1 to 1.4, however, resulted in overall decreased sensitivity, although these effects were most pronounced at low to medium spatial frequencies. Additionally, when *Amp_inh_* was increased, the model was maximally sensitive at higher spatial frequencies when compared with the control, and the sensitivity curve had a more distinct peak. Therefore, *Amp_inh_* had effects on the model sensitivity peak, sensitivity at low to medium spatial frequencies, and preferred spatial frequency.

**Figure 7.**
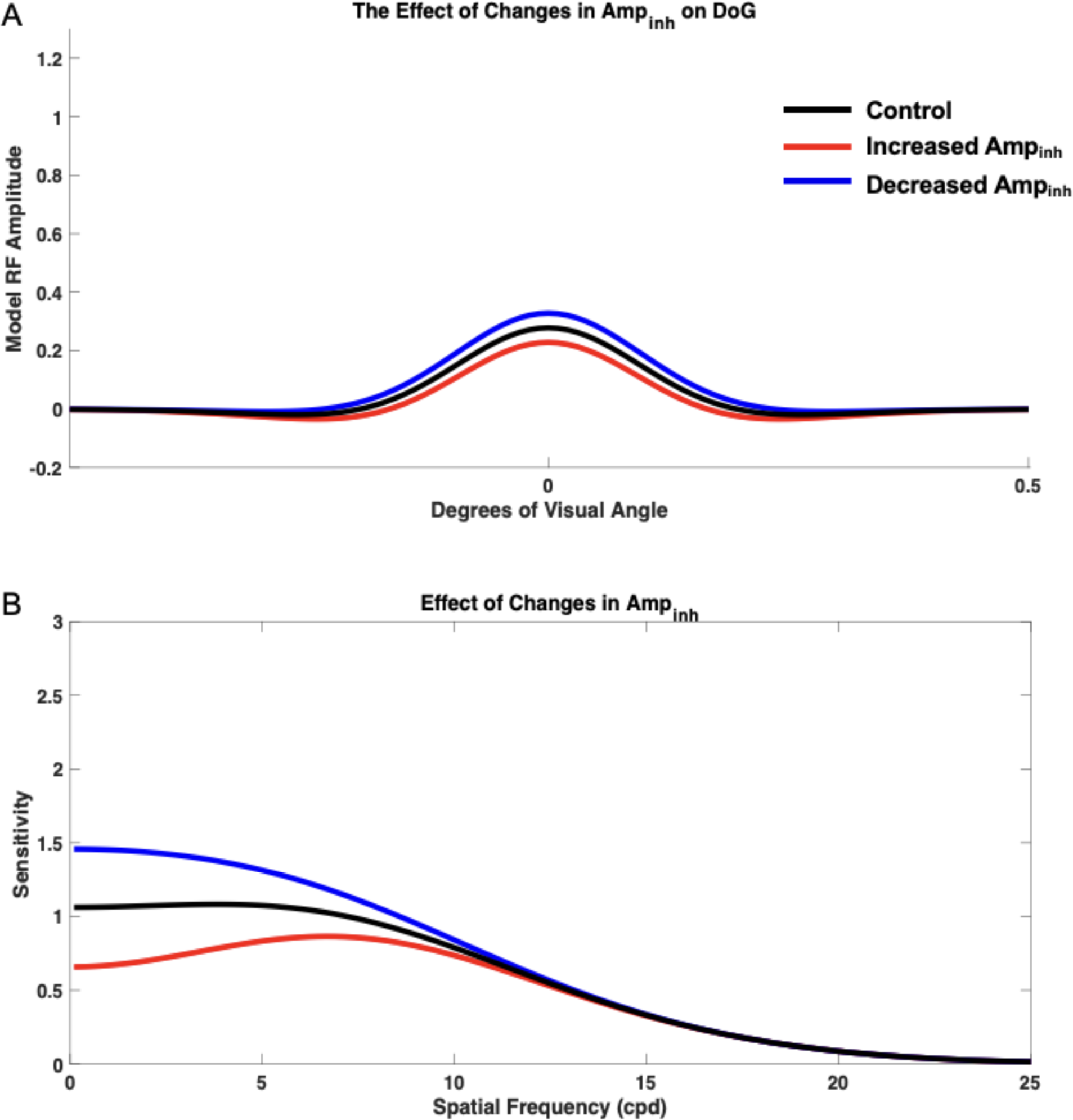
Isolated impact of inhibitory amplitude (*Amp_inh_*) on spatial frequency sensitivity and receptive field profile (DoG). DoG profiles (A) and the impact on spatial frequency sensitivity (B) was compared with the control (black) when *Amp_inh_* was increased to 1.4 (red) and decreased to 0.6 (blue).

#### Impact of Concurrent Changes in Inhibition Extent (σ_inh_) and Amplitude (Amp_inh_)

To determine the impact of concurrent *σ_inh_* and *Amp_inh_* changes, *σ_ex_* and *Amp_ex_* were both held constant at the base (control) value of 1. *Amp_inh_* and *σ_inh_* were then simultaneously varied and the resulting curves were compared with the control sensitivity curve generated with values 1 and 1.6 for *Amp_inh_* and *σ_inh_*, respectively. When *σ_inh_* was decreased to 1.2 and *Amp_inh_* was simultaneously decreased to 0.6, the model demonstrated increased sensitivity at low spatial frequencies, slightly increased sensitivity at medium spatial frequencies, and slightly decreased sensitivity at high spatial frequencies. The most pronounced effect was increased sensitivity at low spatial frequencies (Figure 8 A-B). The model also lacked a distinct peak in the sensitivity curve in this condition. When *σ_inh_* and *Amp_inh_* were both increased to 2, the model demonstrated decreased contrast sensitivity at low spatial frequencies and medium spatial frequencies and slightly increased contrast sensitivity at high spatial frequencies. The effects were most pronounced in the decreased sensitivity to low spatial frequencies (Figure 8 C-D). The model also demonstrated a more distinct peak when compared with the base control curve, and maximum sensitivity occurred at a higher spatial frequency than the control. When *σ_inh_* was decreased to 1.2 and *Amp_inh_* was simultaneously increased to 2, the model demonstrated overall decreased sensitivity across spatial frequencies (Figure 8 E-F). Moreover, the model demonstrated a distinct peak in its spatial frequency sensitivity curve, with maximum sensitivity occurring at higher spatial frequencies than the control condition. When *σ_inh_* was increased to 2 while *Amp_inh_* was decreased to 0.6, the model demonstrated overall increased contrast sensitivity (Figure 8 G-H). In this condition, the model lacked the appearance of a distinct peak in its sensitivity curve. Concurrent variation of *σ_inh_* and *Amp_inh_* demonstrated effects on the appearance of a distinct peak, preferred spatial frequency, and overall contrast sensitivity of the model.

**Figure 8.**
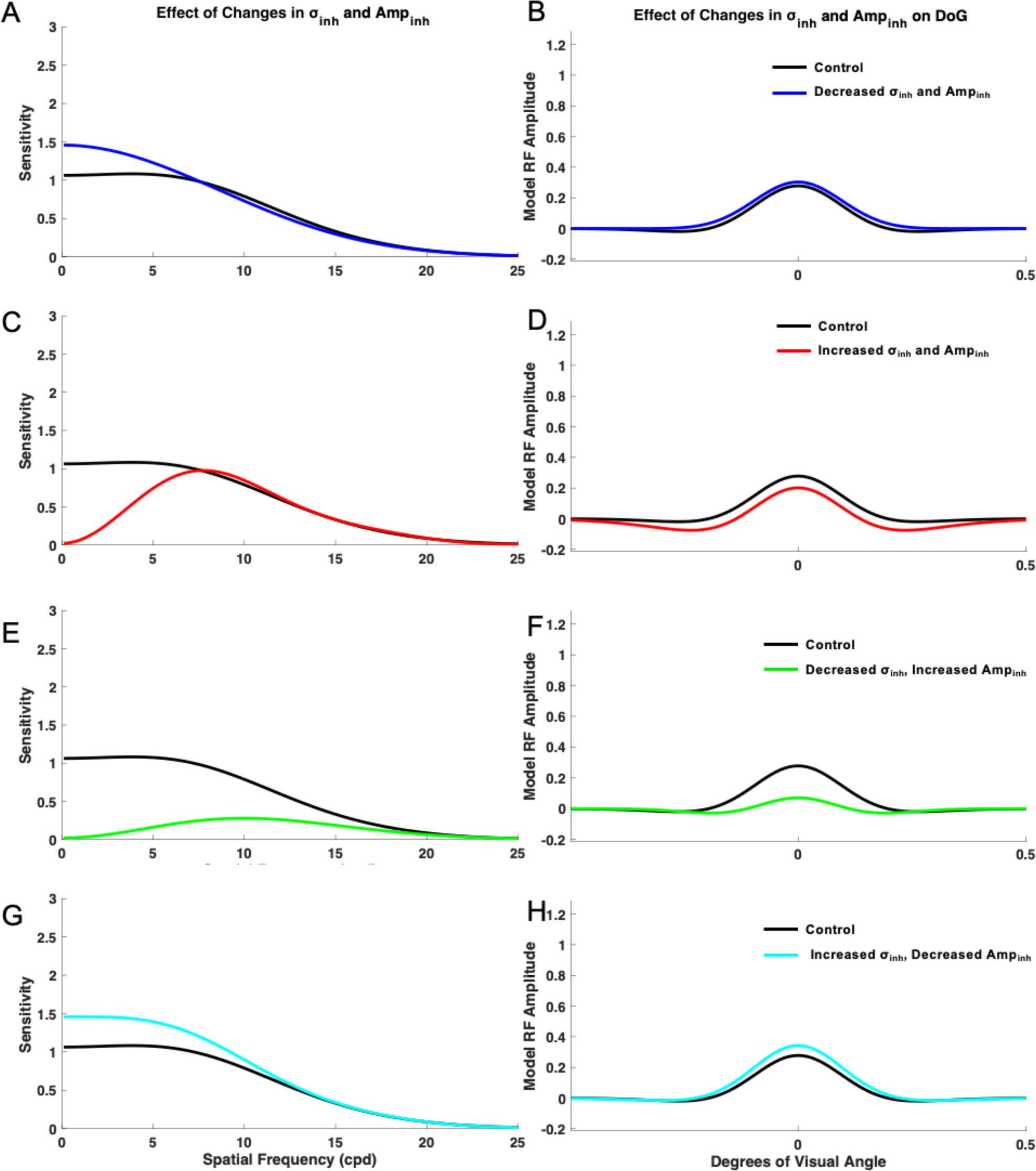
Impact of concurrent changes in inhibition extent (*σ_inh_*) and amplitude (*Amp_inh_*) on the model spatial frequency sensitivity and receptive field profiles (DoG). Impacts were measured when *σ_inh_* was decreased to 1.2 and *Amp_inh_* was decreased to 0.6 (A-B), *σ_inh_* and *Amp_inh_* were increased to 2 (C-D), when *σ_inh_* was decreased to 1.2 and *Amp_inh_* was increased to 2.0 (E-F), and when *σ_inh_* was increased to 2.0 and *Amp_inh_* was decreased to 0.6 (G-H).

### Impact of Changes in Receptive Field Size and Subfields Strength

#### Maintaining the Ratio of Excitation (σ_ex_) to Inhibition (σ_inh_) subfield extents

To determine the impact of the receptive field and its excitatory/inhibitory subfield extents, the raw values of *σ_ex_* and *σ_inh_* were altered, but the base ratio of 1:1.6 was maintained (Figure 9). *Amp_inh_* and *Amp_ex_* were both held at values of 1. To determine the effect of a reduced receptive field size, σ_ex_ and σ_inh_ were both altered by a factor of 0.9. Under this condition, the model showed most pronounced increased sensitivity in the medium spatial frequency range, than in low and high spatial frequencies (Figure 9A-B, blue curve). To determine the effect of an increased receptive field size, *σ_ex_* and *σ_inh_* were both altered by a factor of 1.1. Under this condition, the model demonstrated overall decreased sensitivity (Figure 9A, red curve). In both conditions in which the ratio of *σ_ex_* to *σ_inh_* was maintained at 1:1.6, the contrast sensitivity curves and receptive field profiles demonstrated similar shapes.

**Figure 9.**
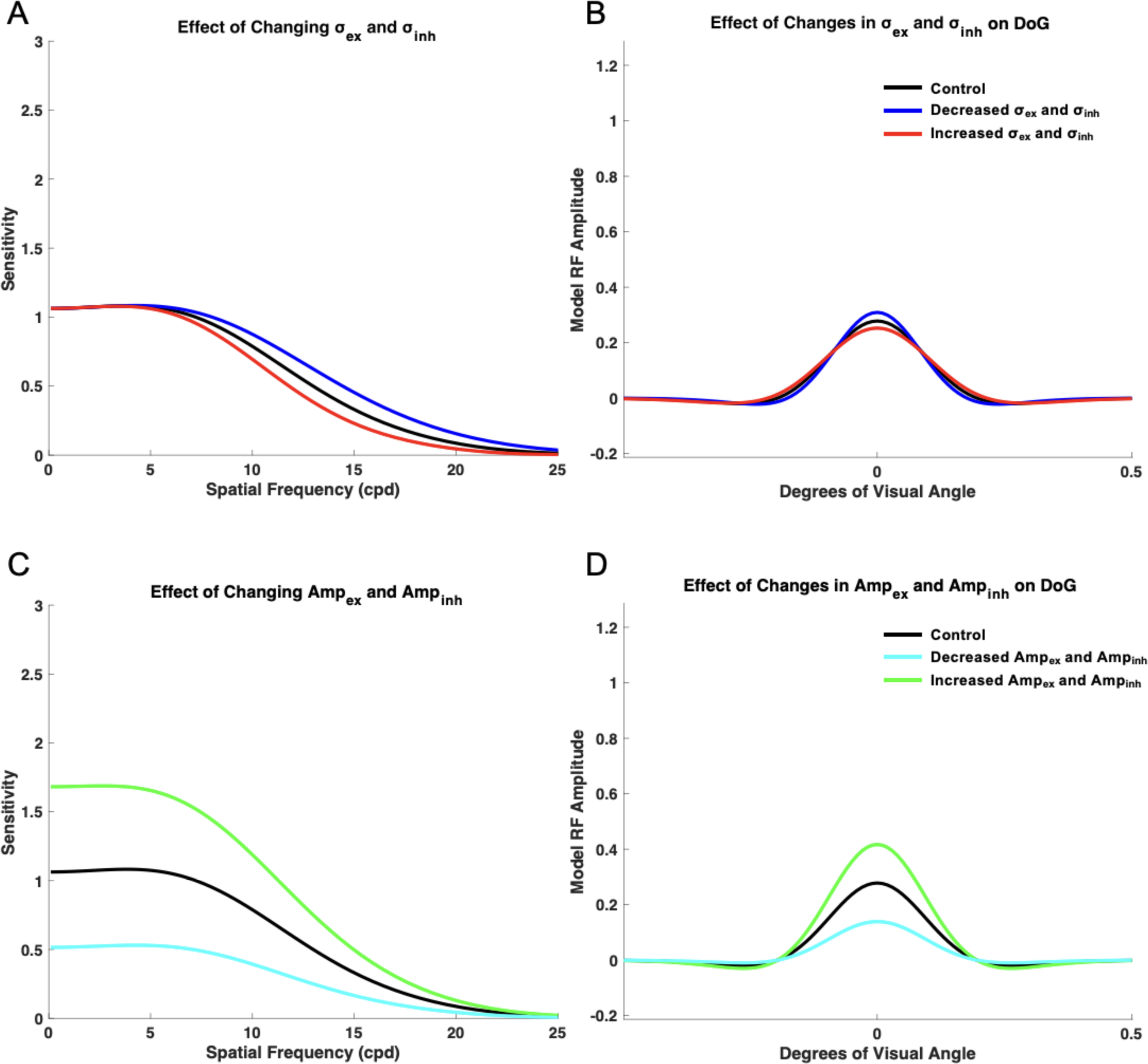
Impact of changing model receptive field size and amplitude while maintaining the ratio of excitatory to inhibitory subfields. Spatial frequency sensitivity (A) and DoG (B); impacts were measured when *σ_ex_* and *σ_inh_* were altered by a factor of 0.9 (blue) and 1.1 (red). Spatial frequency sensitivity (C) and DoG (D); impacts were also measured when *Amp_ex_* and *Amp_inh_* were altered by a factor of 0.5 (cyan) and 1.5 (green).

#### Maintaining the Ratio of Excitatory (Amp_ex_) and Inhibitory (Amp_inh_) Subfields Amplitudes

To determine the effects of the ratio of excitatory to inhibitory subfields amplitude, *σ_ex_* and *σ_inh_* were maintained at base values of 1 and 1.6, respectively. The raw values of *Amp_ex_* and *Amp_inh_* were altered, while keeping the 1:1 ratio of *Amp_ex_*/ *Amp_inh_*. When the values of Amp_ex_ and Amp_inh_ were decreased to 0.5, maintaining the same ratio, the model demonstrated overall decreased sensitivity across spatial frequencies. However, the model did not show a significant shift in the spatial frequency at which maximum sensitivity occurred (Figure 9C-D). When the values of *Amp_ex_* and *Amp_inh_* were increased to 1.5, maintaining the same 1:1 ratio, the model demonstrated overall increased sensitivity across spatial frequencies. The spatial frequency at which maximum sensitivity occurred was also shifted to a lower value (Figure 9C). When the ratio of *Amp_ex_*/ *Amp_inh_* was maintained, the shape of the resulting spatial frequency sensitivity curves resembled that of the base control curve.

## Discussion

In this work, we used a neural model of on-center/off-surround neurons in the early visual system to successfully replicate spatial frequency sensitivity results seen in perceptual data from medicated and unmedicated patients with schizophrenia (Table 3). Based on the model results, we hypothesize that medicated patients may display an increase in the width (spread) and height (strength) of the inhibitory subfield, or either an increase in width (spread) or decrease in amplitude (strength) of the excitatory and inhibitory subfields. Unmedicated patients may display a decrease in the width and amplitude of the inhibitory subfield or a decrease in the width and increase in the amplitude of the excitatory subfield. Below we discuss these findings in the context of the major theories regarding the pathophysiology of schizophrenia and the mechanism of action of common antipsychotic medications. In particular, we correlate our findings with dopaminergic, glutamatergic, and GABAergic abnormalities that have been implicated in the disorder [41–54], the presentation of positive, negative, or cognitive symptoms [42], and the type of antipsychotic medications, which can be classified as either typical, first-generation dopaminergic receptor antagonists, or atypical, second-generation targeting several receptors, [55, 56].

Studies showed that medicated patients with schizophrenia exhibit variable decreases in sensitivity across a wide range of spatial frequencies, showing deficits either at low [3, 7–9], or medium/high [9, 10], or across all frequencies [5]. At low spatial frequencies, increased levels of inhibition best replicated medicated patient perceptual data, in line with evidence supporting increased inhibition in the cortex [57] and impaired inhibitory signaling [58]. Moreover, our model highlighted one plausible mechanism to achieve decreased sensitivity at medium/high spatial frequencies that is in line with the observed reduction of population receptive field sizes in medicated patients with schizophrenia [59]: simultaneous alterations of the excitatory and inhibitory spread with a more dramatic increase in inhibition, so that the ratio of the subfields was not constant. In contrast, multiple parameter combinations resulted in a reduction in the model’s sensitivity across spatial frequencies, in line with behavioral findings and hypotheses based on experimental data [5]. Decreasing the amplitude of excitation and inhibition, which is consistent with studies that have reported reduced interneuron activity and excessive excitatory pruning in schizophrenia [60, 61], provided the best fit. Dampening of the excitatory amplitude, is also supported by new findings that reductions in excitatory synaptic gain, may be linked to the pathophysiology of schizophrenia [62]. Lastly, narrower model inhibition or wider excitation caused imbalance and lower model sensitivity that may reflect altered center-surround interactions in patients with schizophrenia [63].

Unmedicated patients with schizophrenia, on the other hand, exhibit the opposite symptoms, demonstrating increases in spatial frequency sensitivity either at low [11, 12], or across all frequencies [5]. An overall decrease in inhibition, best replicated these findings for low spatial frequencies in the simulations, and an increase in model excitation replicated perceptual data across all frequencies. A reduction in the spread and strength of inhibition would be consistent with reduced inhibitory GABAergic interneuron function due to NMDA receptor dysfunction that could lead to increased excitatory glutamatergic signaling downstream [43, 49]. Interestingly, the model best replicated the increased sensitivity across spatial frequencies in unmedicated patients with schizophrenia when the strength of excitation increased but the spread decreased. Increased cortical excitability in unmedicated patients could reflect an increase in the receptive field’s excitatory amplitude [64], while a decrease in excitatory breadth could reflect pruning of excitatory synapses, hypothesized in schizophrenia [60].

Results from previous studies have indicated the possibility that medication type and dosage may alter spatial frequency sensitivity in patients with schizophrenia, affecting perception [3]. For instance, in medicated patients that exhibited decreased contrast sensitivity at medium and high spatial frequencies, increased antipsychotic dosage was associated with more severe deficits in contrast sensitivity [10]. Medicated and unmedicated patients that were tested at a low spatial frequency (0.5 cpd) did not exhibit significant changes compared to controls, when taking atypical antipsychotics, but demonstrated decreased sensitivity when taking typical medication, or increased sensitivity, if not medicated [6]. Other studies have corroborated the finding that decreased sensitivity is seen in patients taking first-generation antipsychotics [9], but some studies found decreased spatial frequency sensitivity even in patient groups taking mostly second-generation antipsychotics [8]. The model was able to replicate the entire range of potential changes in medicated patients (Figure 10).

**Figure 10.**
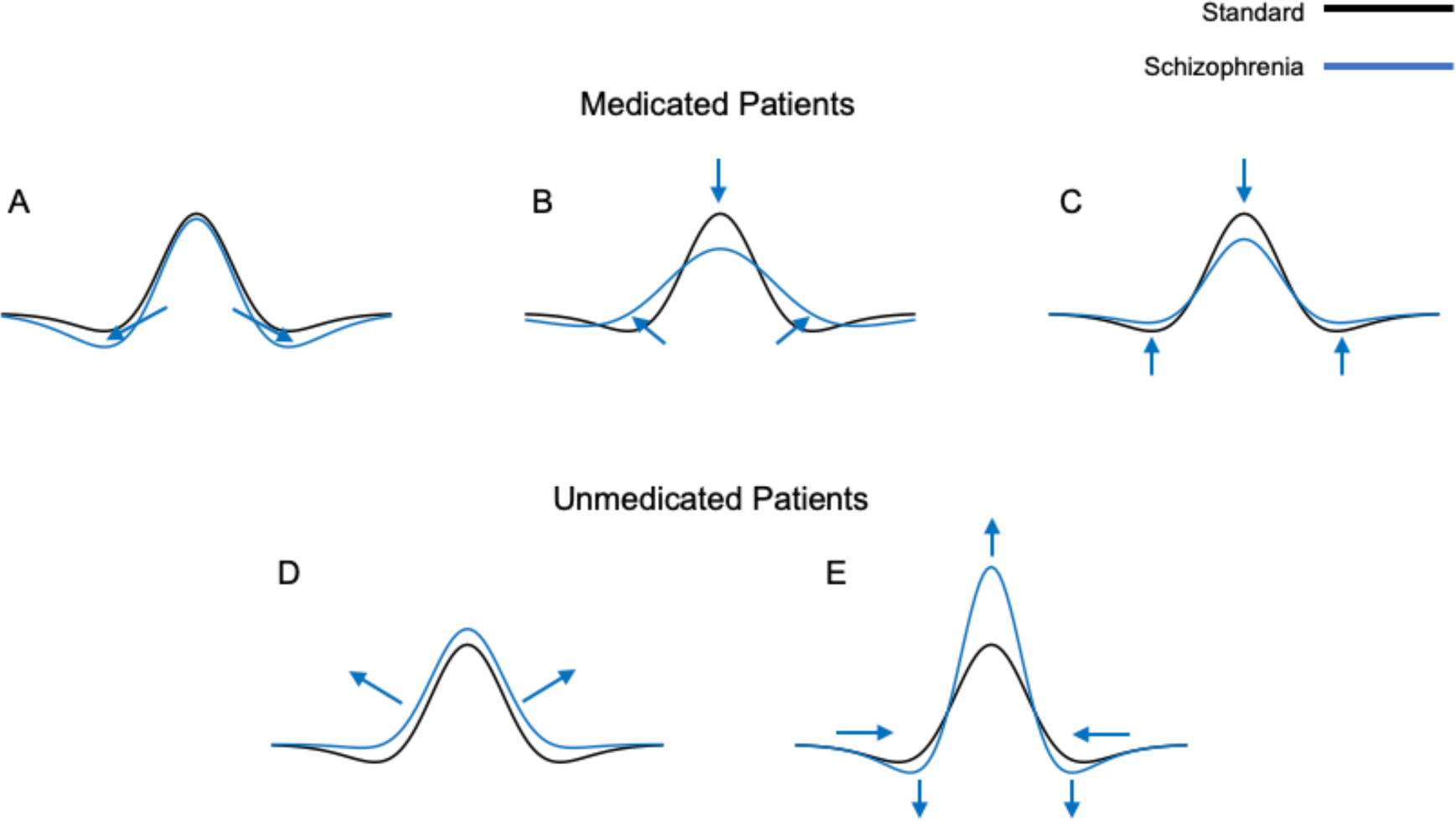
Summary of excitation/inhibition imbalance model parameter settings compared to control to replicate contrast sensitivity of medicated and unmedicated patients with schizophrenia. Based on the existing contrast sensitivity perceptual data in medicated and unmedicated patients with schizophrenia and matching the model’s performance by modifying of its receptive field excitatory and inhibitory subfields, we formulated possible receptive field abnormalities in patients. The formulated model receptive field changes (blue) were compared to the base (control) model receptive field profiles (black). Formulated changes for medicated patients include an increase in the spread (width) and strength (height) of the inhibitory subfield (A), increase in the spread (width) of the excitatory and inhibitory subfields (B), or decrease in the strength (amplitude) of the excitatory and inhibitory subfields (C). Formulated changes for unmedicated patients include decrease in the spread (width) and strength (amplitude) of the inhibitory subfield (D) or decrease in the spread (width) and increase in the strength (amplitude) of the excitatory subfield (E).

Because, first-generation antipsychotics are dopamine D2 receptor antagonists, and their effectiveness is correlated with their D2 receptor binding capacity [55], it is possible that the spatial frequency deficits seen in medicated patients are a result of deficient dopamine signaling. Typical antipsychotics can cause a condition known as drug-induced Parkinsonism, in which dopamine antagonism produces symptoms like those seen in Parkinson’s disease [65]. Patients with Parkinson’s tend to exhibit decreased contrast sensitivity at a range of spatial frequencies, but these effects can be mitigated by drugs such as levodopa, possibly implicating dopamine in contrast sensitivity deficits [66, 67]. Dopamine may also have an effect as early as the retina, where it can weaken gap junctions between horizontal cells, thus reducing receptive field size [47, 68]. In medicated patients, decreased dopamine could lead to an increase in receptive field size and thus decreased spatial frequency sensitivity. If, in unmedicated patients, dopamine signaling is increased at the level of the retina, this could result in a decreased receptive field size and, thus, increased spatial frequency sensitivity.

Thus, if medication type is considered in the context of spatial frequency sensitivity, it is possible that dopamine antagonism produces contrast sensitivity deficits, whereas second-generation antipsychotic medications which have a comparatively lower affinity for dopamine receptors do not produce these same deficits. In addition, antipsychotic medications decrease glutamate metabolite levels over the course of treatment, thus providing a potential insight into the differences observed in spatial frequency sensitivity between medicated and unmedicated patients [45]. Therefore, when comparing perceptual results in medicated patients with those from unmedicated patients, it is important to take note of the medication regimens, as this could present a key variable.

Importantly, NMDA receptors reach their maximum density in the primary visual cortex, where dopaminergic D1, GABA_A_, and GABA_A_/BZ receptors are also particularly dense [69]. It is therefore likely that glutamatergic NMDA receptor hypofunction, GABAergic dysfunction, and dopamine dysregulation in V1 interact to contribute to visual processing abnormalities [48, 53], affecting both unmedicated and medicated patients [70, 71]. Moreover, GABA-related gene expression is altered in patients with schizophrenia [50, 51] and GABA concentration is reduced in V1 of patients with schizophrenia, and this reduction is associated with dysfunction in orientation-specific surround suppression [52, 53].

### Limitations and Simplifications

Our parameterized, rate-based neural model represents a simplification of various early visual areas, such as the primary visual cortex (V1), association visual cortices V2, V4, and the lateral geniculate nucleus (LGN). The single layer model could represent an ensemble of multiple visual areas and their accumulated activities. Since the model is single layer, it is also important to note that it can only approximate network changes that may be occurring in schizophrenia, due to the thalamocortical loop connectivities or those relating to feedback mechanisms [72]. Given that contrast sensitivity is a relatively low-level visual process, this simplification could be a good starting point model for an initial investigation of the underlying mechanisms of related abnormalities seen in schizophrenia [17]. This is relevant given that spatial frequency processing involves both feedforward [73, 74] and recurrent/feedback processing [73, 75]. Although the model does not directly incorporate feedback mechanisms, these elements could be partially underlying broader changes in excitation and inhibition that we examined. With the understanding that the model still represents a simplified version of complicated sensory mechanisms, the model’s “receptive field” is actually a type of population receptive field. This allows changes across different visual areas to be incorporated into a single model layer representing the interconnected areas. This method simplifies the population receptive field properties into *σ* (width, spread) and amplitude (height, strength), thus allowing for the investigation of abnormalities that arise from a variety of detailed neurobiological mechanisms, which impact the inhibitory and excitatory subfields. Therefore, neurotransmitter and receptor-related abnormalities could be simplified, incorporated, and interpreted in terms of changes in excitation and inhibition width/amplitude. Because contrast sensitivity is heavily dependent on the relative balance of the excitatory and inhibitory subfields, this simplified model can provide targeted and valuable insights into contrast sensitivity abnormalities in patients with schizophrenia that arise from changes in one or simultaneous manipulations of several parameters.

### Conclusions

The rate-based, feedforward model we developed demonstrates that the spatial frequency sensitivity abnormalities observed in medicated and unmedicated patients with schizophrenia can be replicated in terms of alterations to the excitatory and inhibitory receptive field subfields. The results indicate that medicated patients with schizophrenia may have increased neural inhibition, altered receptive field size, or concurrently altered levels of excitation and inhibition. Unmedicated patients, on the other hand, may have either increased excitation or decreased inhibition. By utilizing this technique to model and explore the possible pathophysiology of schizophrenia, it is possible to make connections between the hypotheses that exist about this disease and other factors that may influence the observed perceptual deficits.

## Supplementary Materials

**Supplementary Table 1.** Parameter values per model run. Parameters were varied across model trials. Across trials, A was held constant at 1, B was held constant at 10.1, C was held constant at 5, and a was held constant at 0.1. For a given trial, italicized values represent manipulated parameters, and non-italicized values represent values which were held constant. For each parameter, the color of the value matches the color of the respective curve that represents model outcome in Figures 3-9.

| Figure | $Amp_{ex}$ | $Amp_{inh}$ | $\sigma_{ex}$ | $\sigma_{inh}$ |
| --- | --- | --- | --- | --- |
| 3 | 1 | 1 | *0.8, 1.2* | 1.6 |
| 4 | *0.8, 1.2* | 1 | 1 | 1.6 |
| 5A | *0.6* | 1 | *0.6* | 1.6 |
| 5C | *1.4* | 1 | *1.4* | 1.6 |
| 5E | *1.6* | 1 | *0.6* | 1.6 |
| 5G | *0.6* | 1 | *1.4* | 1.6 |
| 6 | 1 | 1 | 1 | *1.2, 2* |
| 7 | 1 | *0.6, 1.4* | 1 | 1.6 |
| 8A | 1 | *0.6* | 1 | *1.2* |
| 8C | 1 | *2* | 1 | *2* |
| 8E | 1 | *0.6* | 1 | *2* |
| 8G | 1 | *2* | 1 | *1.2* |
| 9A | 1 | 1 | *0.9, 1.1* | *1.44, 1.76* |
| 9C | *0.5, 1.5* | *0.5, 1.5* | 1 | 1.6 |

**Supplementary Table 2.**
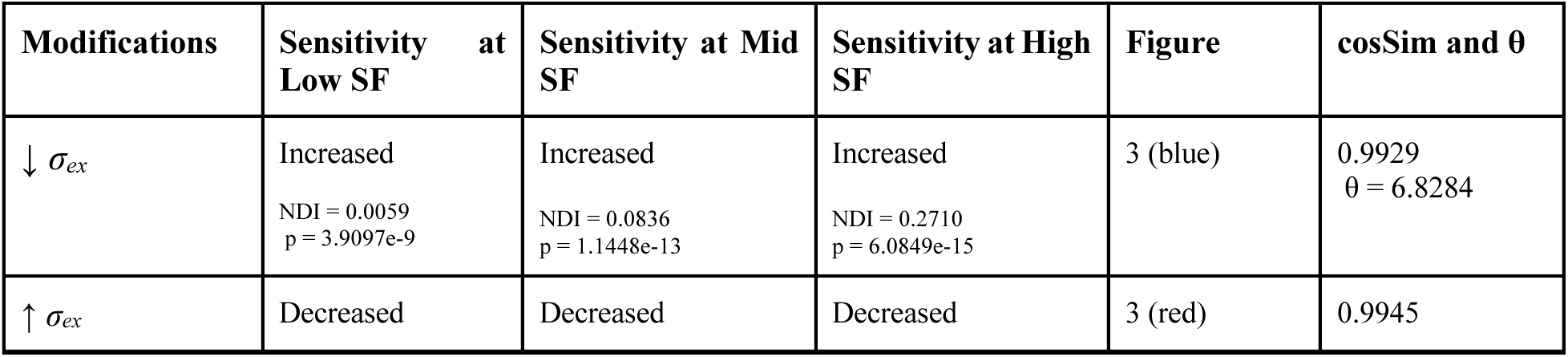
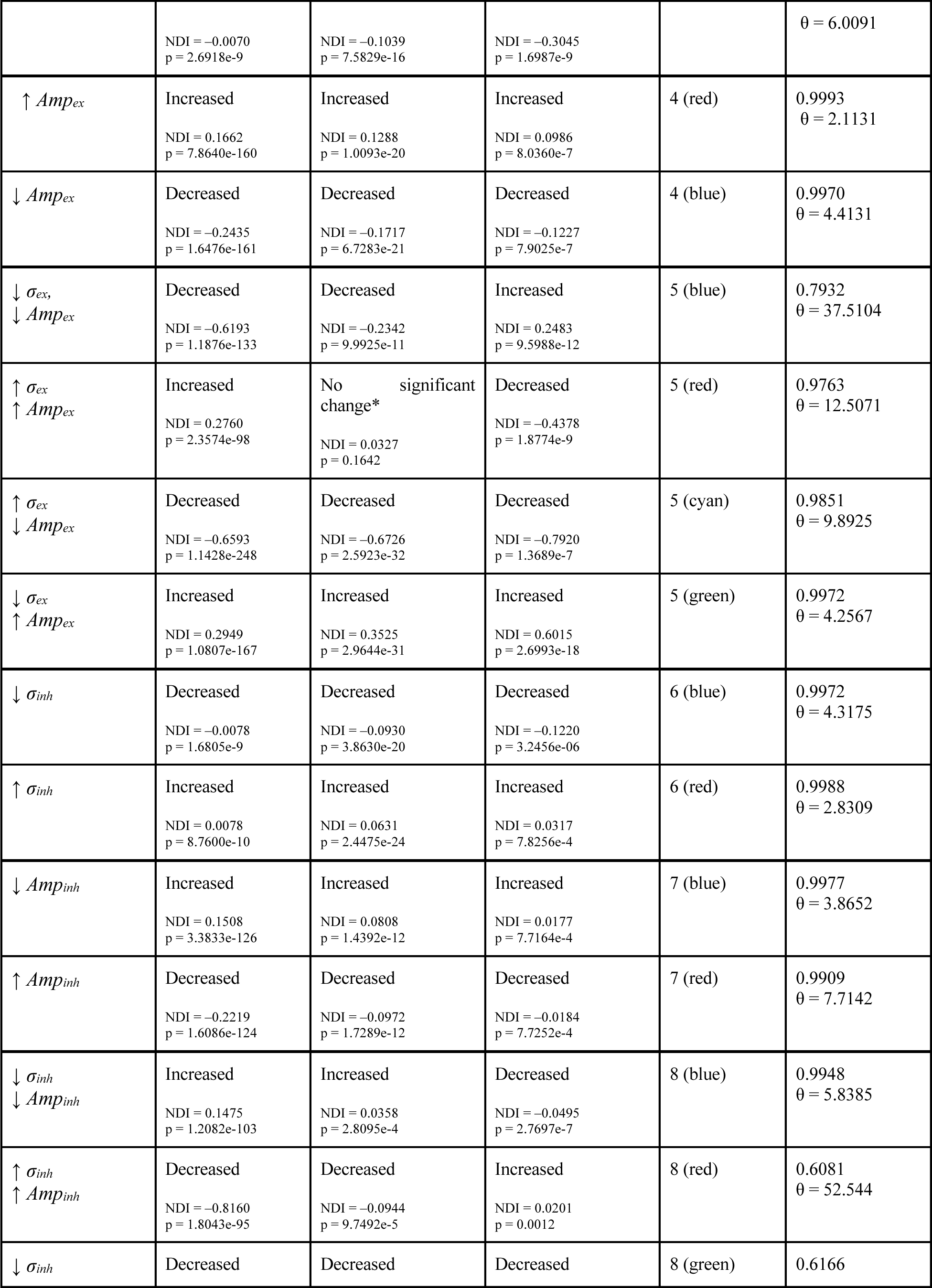
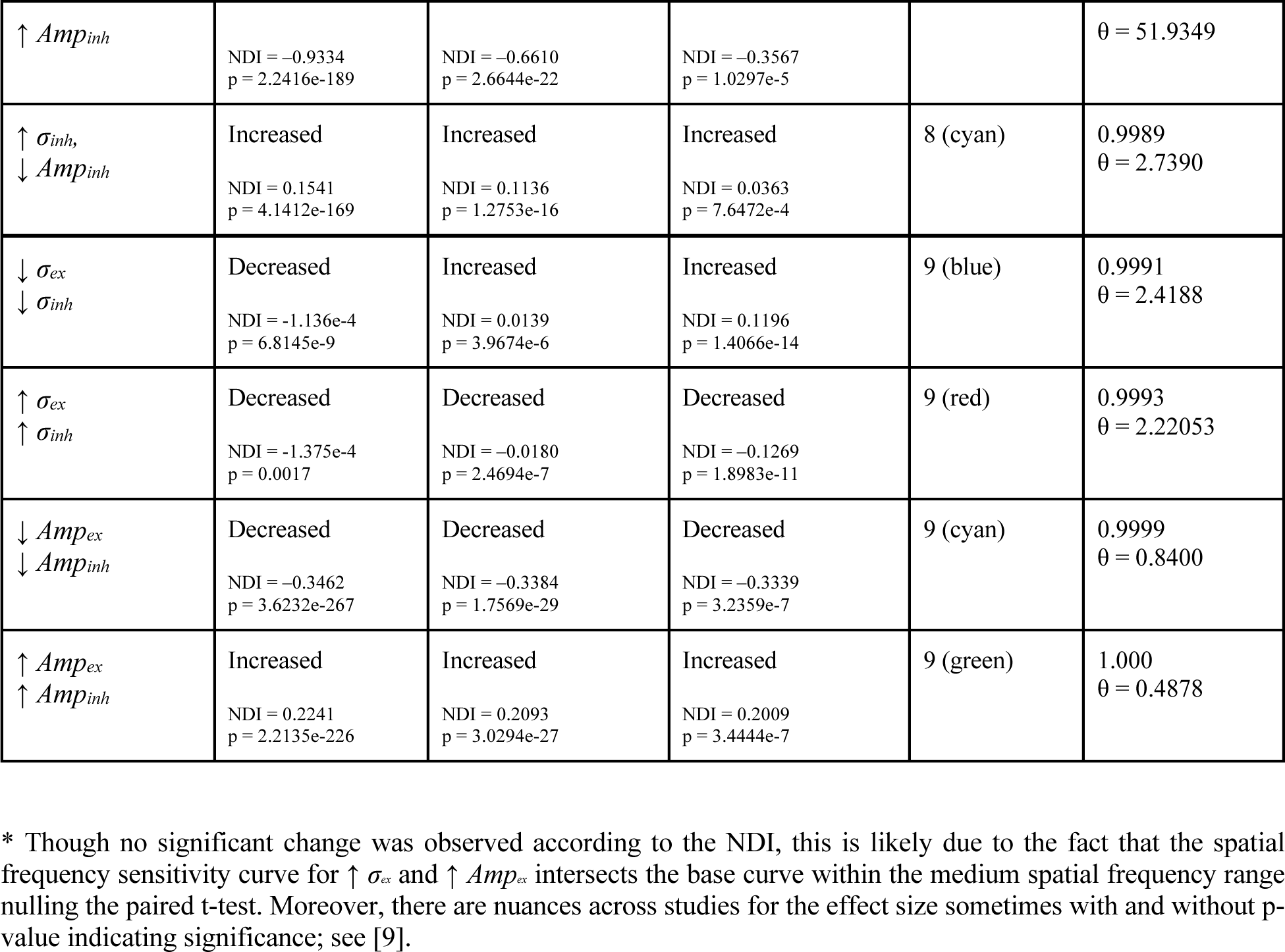
Summary of the model excitation and inhibition parameter search. Comparisons are made based on the model base parameters (control). Positive NDI value indicates increment relative to control, whereas negative NDI value indicates decrement relative to control.

## Notes

### Competing Interest Statement

The authors have declared no competing interest.

